# The stem loop 2 motif is a site of vulnerability for SARS-CoV-2

**DOI:** 10.1101/2020.09.18.304139

**Authors:** Valeria Lulla, Michal P. Wandel, Katarzyna J. Bandyra, Rachel Ulferts, Mary Wu, Tom Dendooven, Xiaofei Yang, Nicole Doyle, Stephanie Oerum, Rupert Beale, Sara M. O’Rourke, Felix Randow, Helena J. Maier, William Scott, Yiliang Ding, Andrew E. Firth, Kotryna Bloznelyte, Ben F. Luisi

**Affiliations:** Department of Pathology, Division of Virology, University of Cambridge, Lab Block Level 5, Addenbrookes Hospital, Hills Road, Cambridge CB2 0QQ, U.K; MRC Laboratory of Molecular Biology, Francis Crick Avenue, Cambridge, CB2 0QH, U.K; Department of Biochemistry, University of Cambridge, Tennis Court Road, Cambridge CB2 1GA, U.K; The Francis Crick Institute, 1 Midland Road, London, NW1 1AT, U.K; Department of Cell and Developmental Biology, John Innes Centre, Norwich Research Park, Norwich, NR4 7UH, U.K; Pirbright Institute, Ash Road, Pirbright, Woking, GU24 ONF, U.K; CNRS-Université Paris Diderot, Institut de Biologie Physico-Chimique, 13 rue Pierre et Marie Curie, 75005 Paris, France; University of California at Santa Cruz, Santa Cruz California, 95064, U.S.A

**Author notes:** These joint-first authors made complementary and equivalent contributions.

**Keywords:** s2m, positive-sense RNA virus, coronavirus, astrovirus, gapmer, LNA, therapeutic oligonucleotides, SARS-CoV-2

## Abstract

RNA structural elements occur in numerous single stranded (+)-sense RNA viruses. The stemloop 2 motif (s2m) is one such element with an unusually high degree of sequence conservation, being found in the 3’ UTR in the genomes of many astroviruses, some picornaviruses and noroviruses, and a variety of coronaviruses, including SARS-CoV and SARS-CoV-2. The evolutionary conservation and its occurrence in all viral subgenomic transcripts implicates a key role of s2m in the viral infection cycle. Our findings indicate that the element, while stably folded, can nonetheless be invaded and remodelled spontaneously by antisense oligonucleotides (ASOs) that initiate pairing in exposed loops and trigger efficient sequence-specific RNA cleavage in reporter assays. ASOs also act to inhibit replication in an astrovirus replicon model system in a sequence-specific, dose-dependent manner and inhibit SARS-CoV-2 infection in cell culture. Our results thus permit us to suggest that the s2m element is a site of vulnerability readily targeted by ASOs, which show promise as anti-viral agents.

## INTRODUCTION

SARS-CoV-2 is a highly infectious virus and the causative agent of the ongoing COVID-19 pandemic. Given the continued rise in cases worldwide, the significant mortality rate and the challenges in predicting the severity of illness in infected individuals (Messner et al., 2020), there is a pressing need for efficacious antiviral therapies (Koirala et al., 2020; Krammer, 2020; https://www.who.int/emergencies/diseases//novel-coronavirus-2019). Moreover, the potential for further outbreaks of infections by emerging pathogenic coronaviruses (Menachery et al, 2015; Akula and McCubrey, 2020) places importance on improving fundamental understanding of coronavirus biology, as well as exploring novel therapeutics to build capability for a rapid response to the next zoonotic jump.

Like other positive-sense (+) single-stranded (ss) RNA viruses, replication of SARS-CoV-2 is orchestrated by virus-encoded enzymes inside infected host cells. The 30 kb long SARS-CoV-2 genomic RNA and the subgenomic mRNA transcripts all contain a common 5’-leader sequence and a common 3’ UTR, which harbour several conserved structural elements, including the stem-loop 2 motif (s2m) (Fig. 1A and B) (Rangan et al., 2020; Kim et al., 2020). The s2m, originally identified in astroviruses (Jonassen et al., 1998), is a highly conserved RNA sequence element, present within the 3’ UTR in the genomes of many astroviruses, some picornaviruses and noroviruses, and a variety of coronaviruses, including members of the subgenus *Sarbecovirus,* which includes SARS-CoV and SARS-CoV-2 viruses (Tengs et al., 2013, Tengs et al., 2016), and *Merbecovirus,* which includes MERS (Frey et al, 2014), the causative agents of three recent severe human pathogenic outbreaks. The SARS-CoV and SARS-CoV-2 s2m sequences are nearly identical, with only 2 point nucleotide divergence (Fig. 1A), in contrast to the overall 20 % genome-wide sequence difference (Kim et al., 2020). The s2m sequence is also highly conserved in the clinical isolates from patients that have tested positive for SARS-CoV-2 during the current pandemic, although with a few isolated exceptions (GSAID database https://www.gisaid.org; UCSC genome browser; Vahed et al.,2020; Yeh and Contreras, 2020). The high degree of s2m sequence conservation is likely to be a direct consequence of a requirement to sustain an elaborate and conserved three-dimensional structure. Indeed, an earlier study of the SARS-CoV s2m element revealed a stable stem loop with a few exposed bases (Fig. 1B) in a 2.7 Å resolution crystal structure (Robertson et al., 2005). Several more recent studies probed RNA accessibility and mapped RNA-RNA interactions of the (+) sense SARS-CoV-2 viral species inside the host cell, confirming that the s2m stem-loop structure folds *in vivo* as well (Huston, 2020; Ziv et al., 2020).

**Figure 1.**
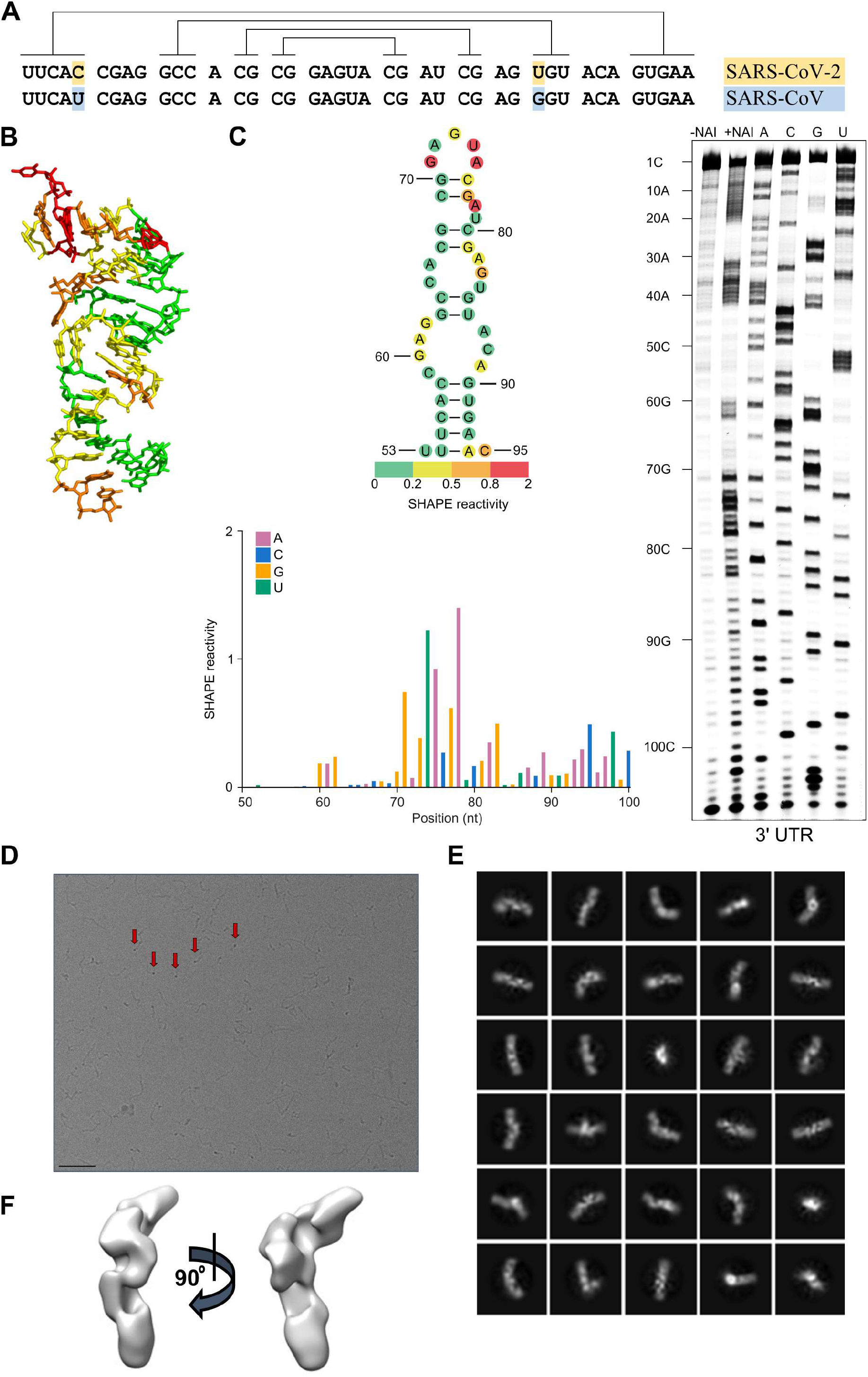
s2m is a conserved structural element in the SARS-CoV-2 genome. (**A**) Sequence alignment of the s2m element in the 3’ UTRs of SARS-CoV-2 and SARS-CoV. Lines indicate base-pairing regions within the element. (**B**) The crystal structure of the SARS-CoV s2m element (adapted from Robertson et al., 2005). (**C**) Chemical probing of the 3’ UTR of SARS-CoV-2. RNA was denatured and refolded in the presence of 100 mM K^+^ and 0.5 mM Mg^2+^, then incubated with NAI (+NAI channel) or DMSO control (-NAI channel). NAI modification was detected by reverse transcription stalling and gel-based analysis. Sequencing lanes were generated by adding ddT (for A), ddG (for C), ddC (for G) and ddA (for U) when performing reverse transcription. The lower panel shows quantification of SHAPE signal in the s2m and flanking regions. Calculation was based on the gel in Fig. 1C, by subtracting the signal of the +NAI lane from that of the -NAI lane. The upper panel shows annotation of SHAPE signal on the s2m structure. The bases with SHAPE signal of 0-0.2, 0.2-0.5, 0.5-0.8 and 0.8-2 were coloured with green, yellow, orange and red, respectively. (**D**) Representative cryoEM image of the SARS-CoV-2 3’ UTR (220 nt) at 2.5 μm defocus. The red arrows indicate features that likely correspond to views along the long axis of duplex regions. The black line in the lower left is a 50 nm scalebar. (**E**) The 2D class averages and (**F**) 3D reconstructions as calculated by CryoSparc 2.15.0.

Because of the apparent high degree of selective pressure to maintain this specific sequence and its structure, the s2m is a promising target for potential antiviral agents, with reduced likelihood of evolving mutations that would lead to resistance. Any agents targeting the s2m element would also have the advantage of acting against a large number of virus RNA species, due to the presence of the element in all (+)-sense virus RNAs, including both the full-length genome and the subgenomic mRNAs. In order to test the accessibility of the s2m element to potential nucleic acid-based therapeutics, we designed a panel of antisense oligonucleotides (ASOs). These oligonucleotides have proven therapeutic potential against viruses and have been undergoing active development for more than a decade (Bennett, 2019). Third-generation ASOs include locked nucleic acids (LNAs), in which a bicyclic linkage at the furanose constrains the conformational freedom of the nucleotide (Singh et al., 1998). LNAs provide high affinity base-pairing to complementary RNA and DNA targets, as well as resistance to nuclease attack (Hagedorn et al., 2018). A version of LNA ASOs known as ‘gapmers’ consists of LNA bases flanking a central DNA sequence (Wahlenstedt et al, 2000). In this design, LNA bases confer resistance to nucleases and provide high-affinity base-pairing to target RNA, while the central DNA region, once base-paired to RNA, recruits ribonuclease H (RNase H), which acts to cleave the RNA in the RNA-DNA duplex. In this process, the DNA is not digested and thus the gapmer remains intact and free to bind further RNA molecules. Gapmers have already been successfully used in clinical trials to silence target transcripts (Bonneau et al., 2019).

In this report, we describe the design and testing of several LNA ASOs (gapmers) against the highly conserved structured s2m element from the 3’ UTR of SARS-CoV-2. We found by chemical probing of the RNA target element that despite the high degree of structural compactness, ASOs were successfully disrupting the s2m structure. Furthermore, gapmers were capable of inducing sequence-specific RNA cleavage *in vitro* and in multiple independent cell-based platforms, including a human reporter system, an astrovirus replicon assay, and direct inhibition of SARS-CoV-2 replication in infected cells. Our results support targeting of the s2m element and other conserved structures with predicted exposed loops in the viral genomes by ASOs. In addition, the particular gapmer designs described here may offer suitable lead compounds for further development as antiviral therapeutics to treat COVID-19 and other diseases caused by RNA viruses possessing the s2m element in their genomes.

## RESULTS

### Model for the s2m element in the context of the SARS-CoV-2 3’ UTR

Given the high sequence similarity between the s2m elements in the SARS-CoV-2 and SARS-CoV genomes (Fig. 1A), the corresponding structures are expected to show high similarity as well. The crystal structure of the SARS-CoV s2m reveals a stem loop with a small pocket that can accommodate cations (Fig. 1B; Robertson et al., 2005) and suggests a similar fold for the SARS-CoV-2 s2m. To test this experimentally, we probed the structure of the SARS-CoV-2 s2m element within the genomic 3’ UTR using SHAPE (**<u>S</u>**elective 2’-**<u>H</u>**ydroxyl **<u>A</u>**cylation analyzed by **<u>P</u>**rimer **<u>E</u>**xtension) (Spitale et al., 2013). The SHAPE reactivity profile strongly agrees with the crystal structure of the SARS-CoV s2m element (Fig. 1C). For instance, high SHAPE reactivities were found at the loop region (G71-A75), indicating strong possibility of a single-stranded nature, while low SHAPE reactivities were found for nucleotides predicted to be base-paired, such as nucleotides 54-58, and nucleotides 90-94. We performed further SHAPE probing on an extended version of the 3’ UTR RNA (“extended 3’ UTR”) that additionally includes ORF10 and the region immediately upstream from it (ORF10 may not be protein coding; Taiaroa et al., 2020; Jungreis et al., 2020) (Fig. S1 and Table 1). We found a high degree of consistency between the SHAPE profiles for the s2m element in the extended and non-extended constructs (Pearson correlation coefficient PCC = 0.996, Fig. S1B and S1C), indicating that the s2m element in the 3’ UTR is stably folded and is unlikely to be affected by flanking regions such as ORF10. Stable structural unit formation by s2m, maintained within an extended surrounding sequence, has also been observed in a recent NMR and DMS chemical probing study (Wacker et al., 2020), with experimental data agreeing with the s2m secondary structure proposed here (Fig. 1C). Further support for our findings comes from recent wholegenome studies of the SARS-CoV-2 RNA structure, which show good agreement between SHAPE reactivity and *in silico* predictions and identify s2m stem loop in this context (Huston, 2020; Manfredonia et al, 2020).

**Table 1.**
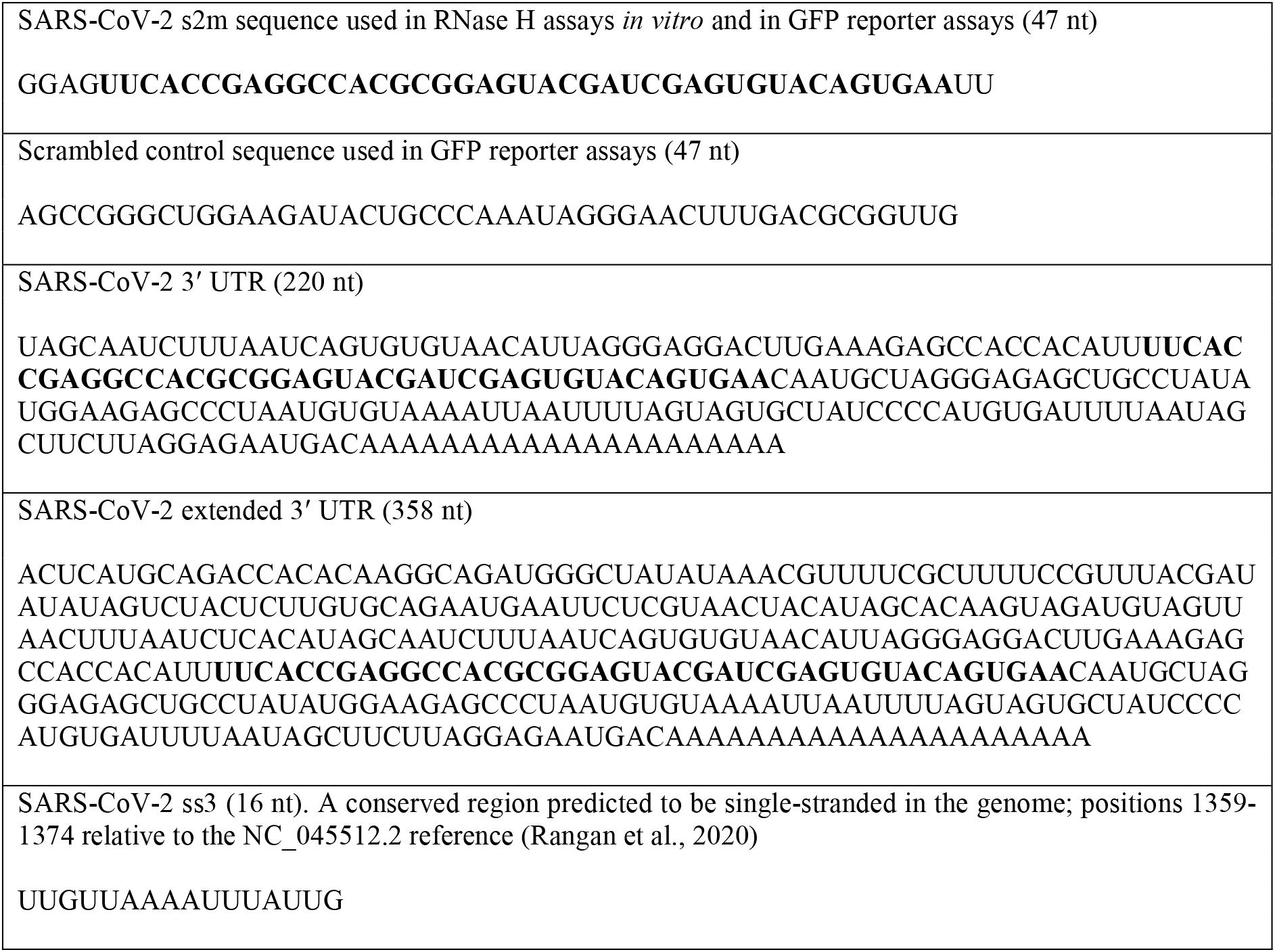
Sequence of the RNA used in this study. The sequence in bold corresponds to the extended s2m region common for these RNA substrates.

To further investigate potential structures formed by the viral 3’ UTR as a whole, we used cryoelectron microscopy. Imaging this RNA fragment mainly yielded elongated shapes resembling thick rope, up to 500 Å in length (Fig. 1D). A subset of the ropey particles were less extended, measuring around 300 Å, which is about half the length expected for an elongated polymer, implying that the RNA there is semi-compact. These particles were subjected to computational 2D/3D averaging in an attempt to reveal underlying shared structural features (Fig 1E and F). However, none could be clearly identified, suggesting that the 3’ UTR RNA does not fold into a well-defined structure, at least *in vitro.*

Our cryo-electron microscopy observations give support to earlier in vivo studies, where cells infected with SARS-CoV-2 virus were probed by cryo-electron tomography (Klein et al., 2020). Both the tomography study and our single-particle imaging reveal small, high-contrast features at the periphery of the RNA particles, which might represent views down the long axis of some duplex RNA regions (Fig. 1D, red arrows).

Overall, based on the prediction from the SARS-CoV s2m RNA crystal structure and our SHAPE probing results, we conclude that the s2m element in the 3’ UTR of SARS-CoV-2 RNA folds into a stem-loop structure, which may be highly conserved among coronaviruses. This is in contrast to the entire 3’ UTR as a whole, for which we do not observe any well-defined global structure.

### Design of LNA ASO gapmers against the s2m element; assessment of binding and activity *in vitro*

Although the highly conserved nature of the s2m element makes it an attractive target for therapies based on ASOs, the structured nature of the target may potentially interfere with ASO-target base-pairing. To facilitate gapmer-induced disruption of the native s2m structure, we designed gapmers so that the high-affinity LNA bases would pair with the RNA bases predicted to be exposed in the s2m element (Fig. 2 and Table 3). This pairing should facilitate initial gapmer-target interaction, hypothetically leading to unfolding of the s2m element as the rest of the gapmer base-pairs with the complementary target nucleotides.

**Figure 2.**
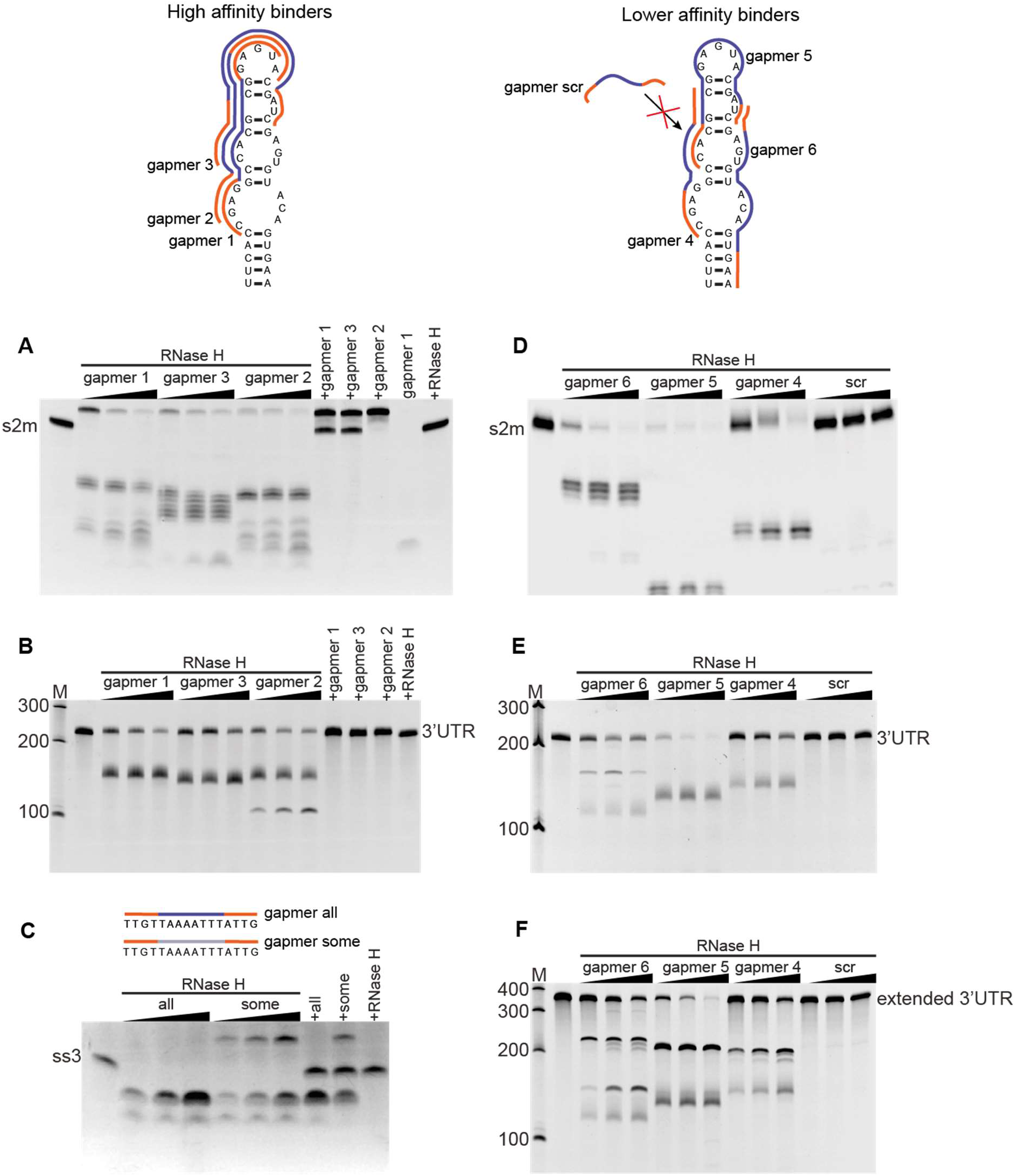
Antisense oligomers direct RNase H cleavage of the s2m element and a conserved single stranded region (ss3) in vitro. The upper panel shows the design of the six gapmers that are complementary to the s2m used in this study, as well as a non-specific control gapmer scr (Table 3). The LNA is indicated in orange, phosphorothioate-linked DNA in blue, phosphodiester-linked DNA in light grey. RNase H cleavage of the isolated s2m (**A, D**), 3’ UTR (**B, E**), the extended 3’ UTR (**F**) and the predicted single-stranded region ss3 (**C**). Three target to gapmer molar ratios were tested: 1:05, 1:1 and 1:2. Incubation of RNA target with RNase H alone does not lead to cleavage (+RNase H, last lane), and is not driven by control gapmers with scrambled DNA sequence (scr). Incubation of RNA target with gapmer without the addition of RNase H does not lead to degradation either, but does lead to the appearance of a retarded band that likely corresponds to target:gapmer duplex.

**Table 2.**
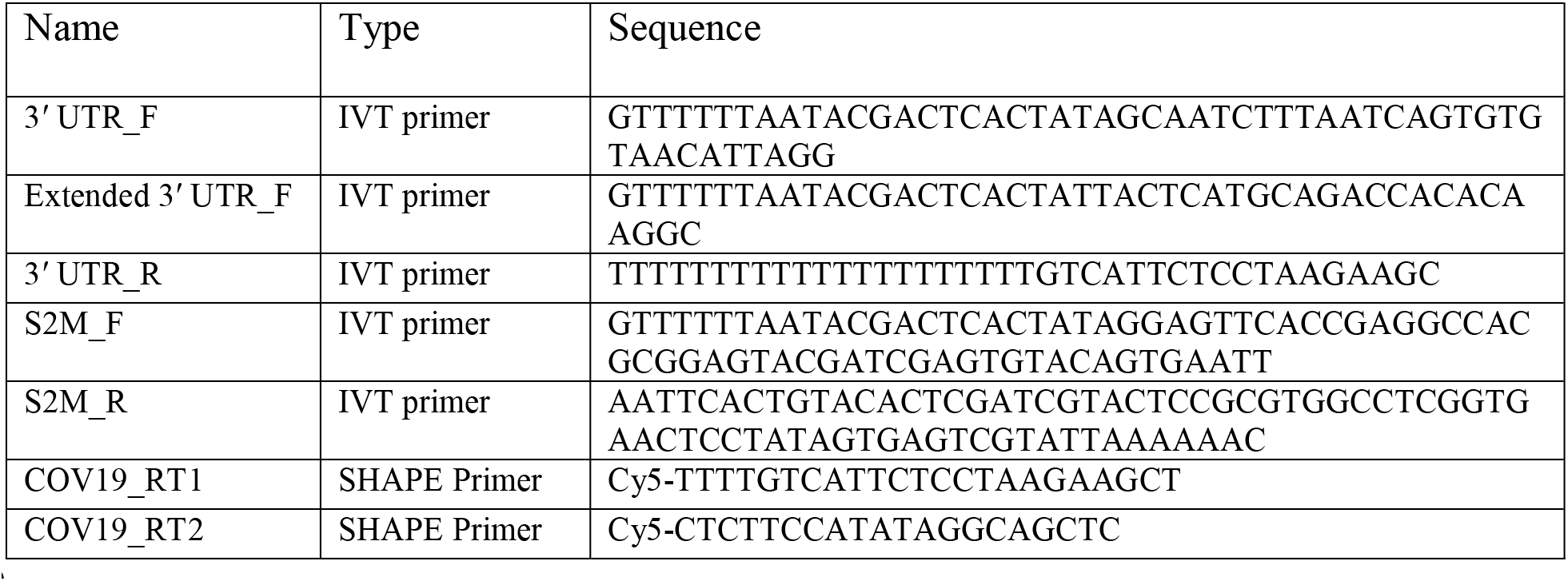
Primers for IVT and SHAPE analysis

**Table 3.**
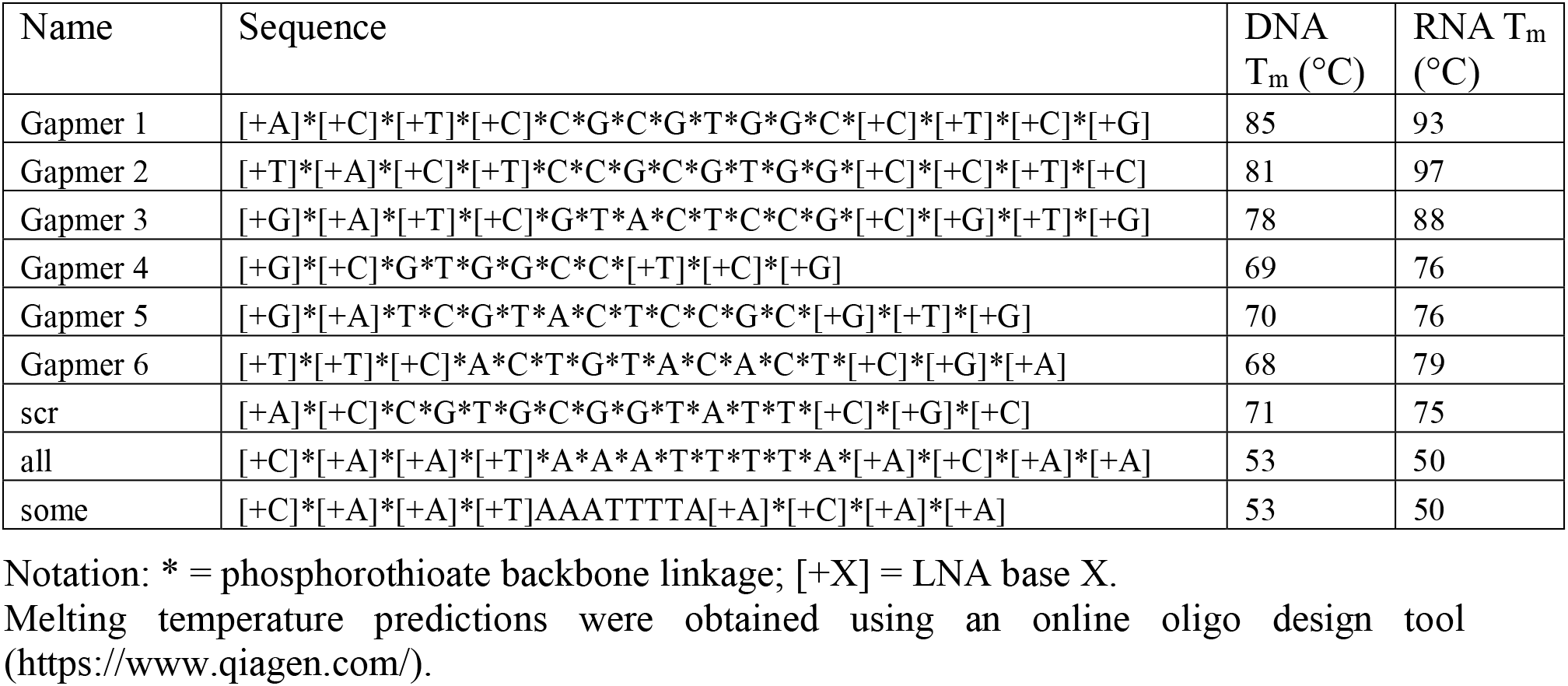
Gapmers used in this study.

We designed a panel of gapmers and tested their ability to direct RNase H cleavage *in vitro*, using s2m RNA as the substrate for the purified enzyme (Fig. 2A and D). Gapmers 1-3 were designed to have higher affinity for target RNA than gapmers 4-6, as indicated by respective predicted melting temperatures (schematic top panel, Fig. 2 and Table 3). The presence of the sequence-specific gapmers 1-6 in the digestion reactions led to clear degradation of target RNA, regardless of their predicted melting temperature bracket, whereas non-sequencespecific control gapmer with scrambled sequence failed to have an effect (“scr”, Fig. 2D). The degradation was very efficient even for 2:1 s2m:gapmer molar ratio, indicating that gapmers can be recycled and can direct multiple turnover of substrates by RNase H. Gapmers 1-6 also drove cleavage of the whole 3’ UTR (Fig. 2B and E) and of the “extended 3’ UTR” construct that additionally includes ORF10 and the region immediately upstream of it (Fig. 2F), indicating that the target s2m sequence is successfully recognised and is accessible for gapmer base-pairing in its native sequence context. The observation that both the higher-affinity gapmers 1-3 (Fig. 2A and B) and the lower-affinity gapmers 4-6 (Fig. 2 D, E and F) were able to direct RNase H cleavage of the s2m element indicates that a range of gapmer-target affinities are compatible with successful target degradation of the highly structured s2m sequence. This flexibility in gapmer design is particularly important as higher affinity gapmers may be expected to have more off-target interactions in cells, which may be reduced in lower-affinity variants.

Additionally, we designed and tested gapmers targeting a single-stranded conserved region in SARS-CoV-2, at position 1359-1374 relative to the NC_045512.2 virus reference genome (“ss3”; Rangan et al., 2020) and tested alternative gapmer backbone chemistries with these. Gapmers designated “all” and “some” have the same sequence and base composition, but different polymer backbones (Table 3). The entire backbone of gapmer “all” contains phosphorothioate modifications, as is also the case in gapmers 1-6; it is a well-established modification conferring some nuclease resistance to oligonucleotides (Kumar et al., 1998; Eckstein, 2000). The backbone in gapmer “some”, however, is mixed; DNA bases are linked by phosphodiester backbone, whereas LNA bases are linked by phosphorothioate backbone.

*In vitro* digestion experiments (Fig. 2C) indicate that both chemistries are compatible with RNase H recruitment and target RNA degradation and also suggest general applicability of gapmer-induced degradation of viral RNA sequences. Gapmer “some” appears to have higher binding affinity for the “ss3” target, judging from the presence of an extra band at the top of the gel in Fig. 2C, which likely corresponds to a gapmer-target dimer. This is consistent with expectation, as phosphorothioate modifications are known to reduce target affinity (Grünweller et al., 2003), so gapmer “all” would be expected to show weaker binding than gapmer “some”. On the other hand, gapmer “all” generates a higher amount of cleaved product (Fig. 2C), which supports our choice of phosphorothioate backbone throughout gapmers 1-6.

### s2m structure is successfully remodelled by LNA gapmers

To test whether gapmers have any effect on the s2m structure, we performed SHAPE probing of SARS-CoV-2 3’ UTR in the presence of gapmers 1, 2 and 3 targeting s2m and gapmer “all” as a non-specific control. Since SHAPE probing detects the accessibility of nucleotides (Spitale et al., 2013), it is capable of revealing both intra- and inter-molecular RNA base-pairing interactions.

In the presence of gapmer 1 and gapmer 2, SHAPE reactivity profiles strongly decreased in the regions targeted by the gapmers, indicating inter-molecular interactions formed between gapmers and their target sequences (Fig. 3A-D). In contrast, SHAPE reactivities strongly increased in the regions that were originally base-paired with the gapmer-target regions and in their flanking regions, indicating they were more single-stranded in the presence of gapmers, consistent with expectation.

**Figure 3.**
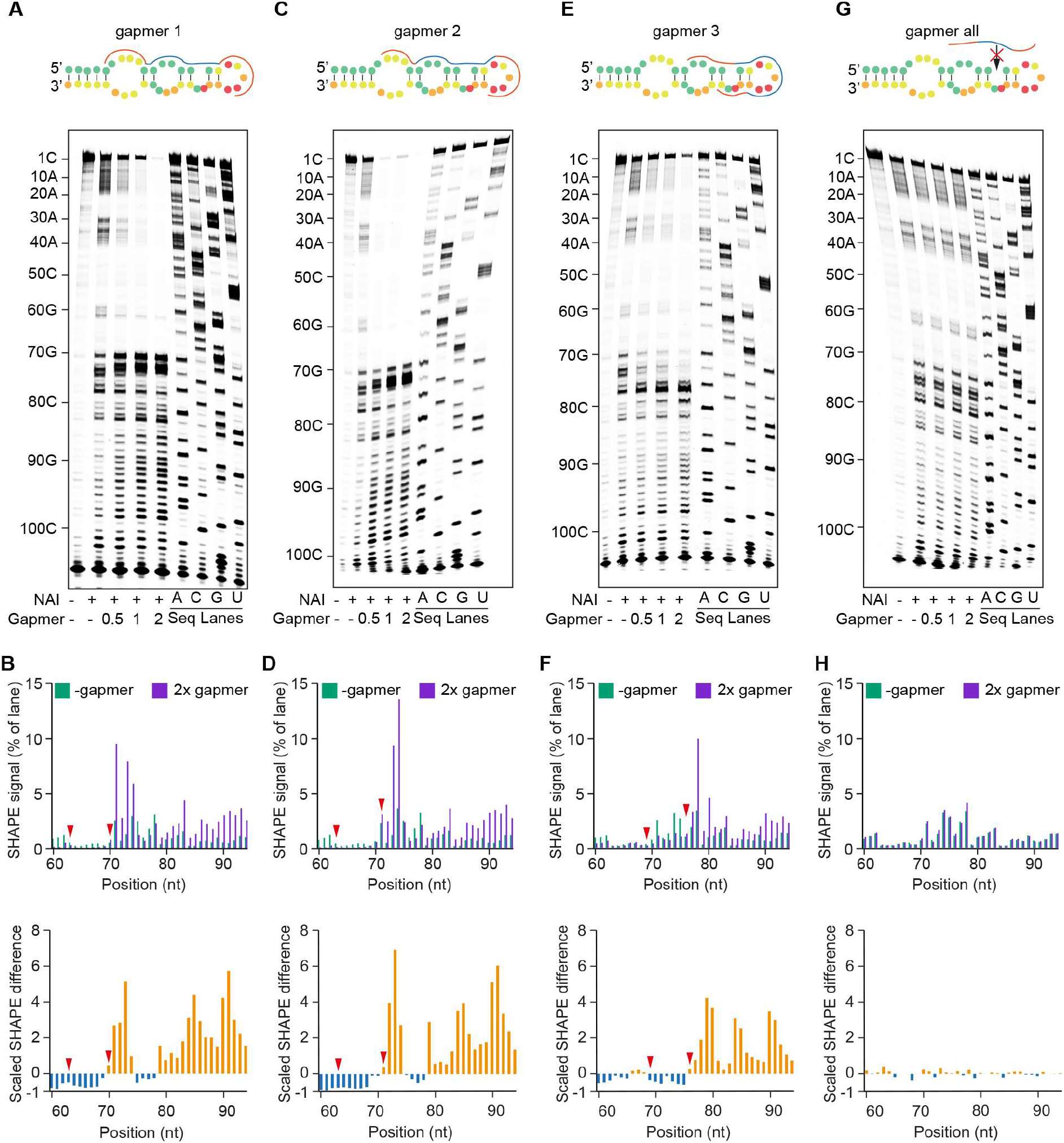
SHAPE probing reveals RNA structure changes induced by LNA gapmers. (**A, C, E, G**) SHAPE probing of SARS-CoV-2 3’ UTR structure in the presence or absence of the gapmer indicated. RNA was denatured and refolded in the presence of 100 mM K^+^ and 0.5 mM Mg^2+^, then incubated with different amounts of gapmer (0×, 0.5×, 1×, 2× that of RNA) and probed using NAI. (**B, D, F, H**) Quantification of A, C, E and G, respectively. Analysis of the differences in SHAPE signal from SARS-CoV-2 3’ UTR alone and in the presence of 2× gapmer. Red arrows indicate the start and end points of gapmer target regions. (**A** and **B**) The presence of gapmer 1 induced an increase in SHAPE signal at positions 70-74 and 79-94, highlighted in orange, indicating that these nucleotides are more unstructured. A strong decrease in SHAPE signal was observed at positions 60-69, highlighted in blue, indicating decreased accessibility of these bases, which could be caused by their base-pairing with the gapmer. (**C** and **D**) The reactivity profile in **D** is similar to that in **B**, due to the similar target regions of gapmer 1 and gapmer 2. (**E** and **F**) In the presence of gapmer 3, nucleotides at positions 69-75 are more structured, while nucleotides at positions 76-94 are less structured, as indicated. **(G** and **H**) No significant differences in SHAPE signal could be detected in the presence or absence of the non-specific control gapmer “all”, indicating that it is unable to cause structural changes in the SARS-CoV-2 3’ UTR.

In the presence of gapmer 3, which targets the single-stranded loop region within s2m, SHAPE reactivity profile showed a strong decrease in this region (Fig. 3E and F), indicating successful targeting by gapmer. We also found that this inter-molecular interaction between gapmer 3 and the loop region led to increased SHAPE reactivities downstream of the loop region. This suggests that inter-molecular interactions between gapmers and target regions could also remodel the folding status of flanking regions.

Notably, the observed changes in SHAPE reactivity were dependent on the concentration of gapmers and were not detected in the presence of a control gapmer (gapmer “all”) which is not able to target the s2m element (Fig. 3G and H). Taken together, our results validate the design of gapmers in targeting the s2m element and highlight the effect of gapmers in remodelling s2m structure.

### s2m acts to direct gapmer-induced reporter gene silencing in human cells

To investigate if gapmers against the s2m element could drive target RNA silencing in human cells, we set up a tissue culture-based reporter system. We generated lung-derived A549 and HeLa cell reporter lines carrying stably integrated GFP genes that encode either the wild type s2m element or a scrambled control sequence in the 3’ UTR (Fig. S2A and B). Transfection with gapmers against s2m reduced GFP fluorescence levels in the two s2m-containing cell lines for both the higher-affinity gapmers 1-3 and the lower-affinity gapmers 5-6 (Fig. 4A and B). Gapmer 4, the shortest of the tested gapmers and containing only six DNA nucleotides (Table 3), had no effect (Fig. 4A and B); this may reflect difficulties in recruiting human RNase H in vivo to this length of RNA-DNA duplex (Kurreck et al., 2002). Non-specific control gapmers did not affect GFP fluorescence levels, nor did treatment with s2m-specific gapmers of control cell lines in which the sequence of the s2m element was scrambled (Fig. S3A and B). These results indicate that the silencing effect relies on sequence-specific gapmer-target interaction. Overall, the results show that gapmers against the s2m element have the potential to silence gene expression from mRNAs containing s2m in their 3’ UTR sequences, which is the case for SARS-CoV-2 mRNAs.

**Figure 4.**
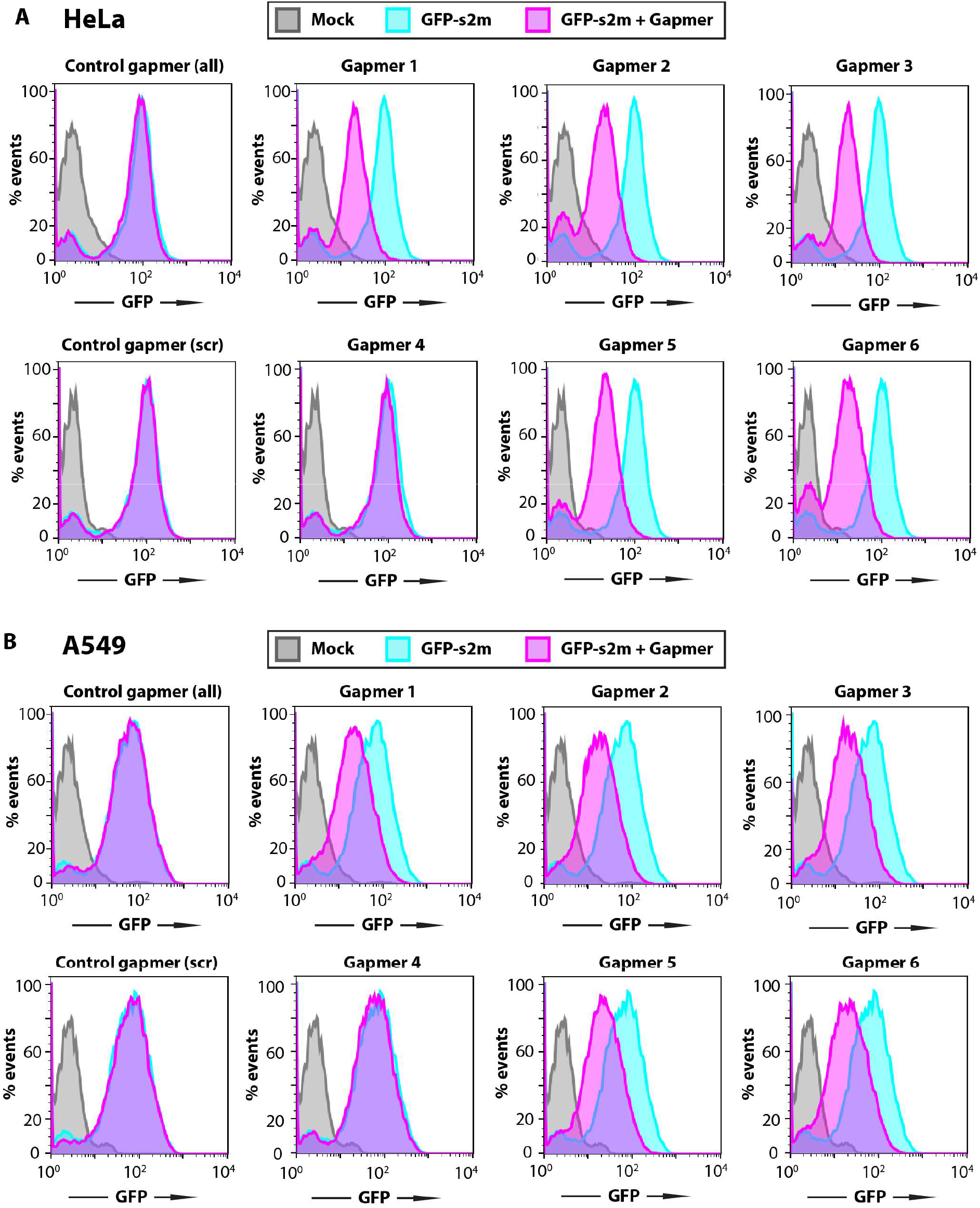
Gapmer-induced reduction of protein levels in cell reporter assays. Flow cytometry analysis of the GFP expressing cells. HeLa (**A**) and A549 (**B**) cell lines containing a genomic insertion of a GFP reporter construct with the s2m sequence in its 3’ UTR (GFP-s2m) were transfected with 20 nM of the indicated gapmers and analysed 72 h post-transfection by flow cytometry. Treatment with gapmers against the s2m element, but not a nonspecific control gapmer, induced reduction in fluorescence, as detected by flow cytometry. Data are representative of three independent experiments.

Our experimental method also functioned to test the hypothesis that the s2m is a post-transcriptional response element. We used the GFP reporter cell lines to compare fluorescence levels between the GFP-s2m and GFP-scrambled cells. No difference was observed, indicating that the s2m element itself does not affect fluorescent protein production (Fig. S2C) suggesting that it does not act as an independent element in *cis* in translation of viral mRNAs.

### LNA gapmers against s2m inhibit replication in an astrovirus replicon model system in human cells

To investigate the effects of s2m-targeting LNA gapmers on viral replication, initially we employed an astrovirus replicon system. Many (+) ssRNA viruses share a similar repertoire of genetic elements required for the replication of viral RNA, and this is true in the case of coronaviruses and astroviruses. Despite large differences in genome size, coronaviruses and astroviruses possess a similar modular organization, including the order of non-structural and structural genes, a frameshift signal to access the RNA-dependent RNA polymerase (RdRp) open reading frame, and production of 3’-coterminal subgenomic mRNAs for structural and accessory protein expression. Like coronaviruses, many astroviruses - including human astrovirus 1 (HAstV1) - contain an s2m element in the 3’ UTR of their genomes (Fig. 5A). We have recently developed a robust HAstV1-based replicon system (Fig. 5A, lower panel), which permits the evaluation of RNA replication in multiple cell types (Lulla and Firth, 2020). The small astroviral genome size (~7 kb) compared to coronaviruses (~30 kb), allows for rapid manipulation of sequences for anti-viral testing in a less restrictive environment than that required to manipulate SARS-CoV-2.

**Figure 5.**
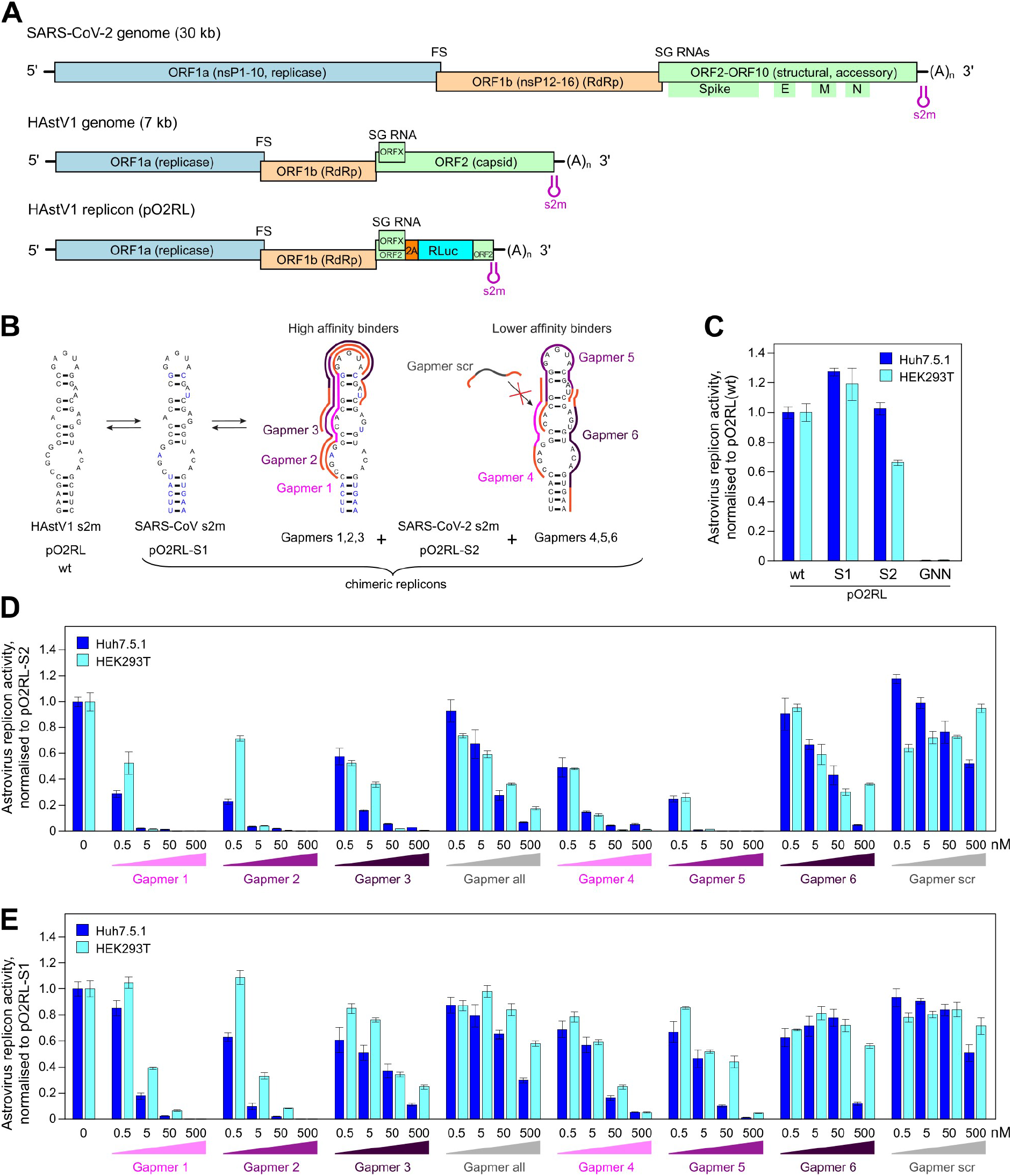
Inhibition of astrovirus replicon activity by gapmers targeting the SARS-CoV-2 s2m RNA element. (**A**) Schematic of the SARS-CoV-2 and human astrovirus 1 (HAstV1) genome organisation. The lower panel represents the astrovirus replicon (pO2RL). FS, frameshift signal; SG, subgenomic; RLuc, Renilla luciferase; RdRp, RNA dependent RNA polymerase. The presented virus and replicon genomes are not to scale. (**B**) Conservation of the s2m 3’ UTR element (two-dimensional representation) between HAstV1, SARS-CoV and SARS-CoV-2. In the astrovirus replicon, the HAstV 1 s2m was switched for the SARS-CoV or SARS-CoV-2 s2m; the wild-type and chimeric replicons are indicated below. Gapmers (1, 2, 3 and 4, 5, 6) are colour-coded in light, medium and dark magenta, respectively. (**C**) Luciferase activity of the wildtype, chimeric, and replication-deficient (RdRp GDD motif mutated to GNN) astrovirus replicons measured in Huh7.5.1 (dark blue bars) and HEK293T (light blue bars) cells. (**D**) Inhibition of the SARS-CoV-2 chimeric replicon by gapmers at 0.5-500 nM concentration range. (**E**) Inhibition of the SARS-CoV chimeric replicon by gapmers at 0.5-500 nM concentration range. For D-E, all data are presented as mean ± s.d.; *n* = 3 biologically independent experiments, full data and statistical analyses are provided in Supplementary Tables S1 and S2. Replicon activity is presented as the ratio of Renilla (subgenomic reporter) to Firefly (cap-dependent translation, loading control), normalized by the same ratio for the untreated control replicon.

Building on promising results in a cell-based reporter system, we used the astrovirus system to test gapmer efficacy in the context of virus-like replication, where replicative intermediates are generally physically occluded within host membrane derived vesicles, as is also the case in *bona fide* virus infection. We generated chimeric astrovirus replicons bearing the s2m elements from SARS-CoV or SARS-CoV-2 (Fig. 5B). Chimeric replicons recapitulated the replication properties of the wild-type astrovirus replicon (Fig. 5C), indicating that this system is suitable for testing gapmers against multiple s2m sequences. To rule out any potential cell-specific effects, all gapmers were tested in two human cell lines - Huh7.5.1 (Lulla and Firth, 2020) and HEK293T (optimised for this study). The replication of replicons bearing SARS-CoV-2 s2m sequences was efficiently inhibited by gapmers 1, 2, and 5, causing inhibition in the subnanomolar range, with a less pronounced effect found for gapmers 3, 4, and 6. The inhibition of non-specific control gapmers (“all” and “scr”) was significantly below their composition-matched counterparts, gapmers 1-3 and 4-6, respectively (Fig. 5D). The same gapmers were also tested against the SARS-CoV s2m in this system and found to be active, though with a lower potency (Fig. 5E). This could potentially be attributed to differences arising from C-G versus C-U juxtaposition within the respective s2m elements (Fig. 5B), leading to changes in the s2m structure (Aldhumani et al., 2021), thus potentially affecting the s2m gapmer binding properties. Replication in the presence of sufficient concentrations of gapmers 1, 2 or 5 dropped to the baseline level of the pO2RL-GNN mutant, which is completely deficient in replication due to a mutated RdRp active site (Fig. 5C).

To assess the cytotoxicity and potential off-target effects of the tested gapmers, we (i) performed a lactate dehydrogenase release-based cytotoxicity assay, and (ii) evaluated the efficiency of cap-dependent translation in the presence of the different gapmer concentrations. Consistent with previous work (Kaur et al., 2007), these assays showed no gapmer-induced cellular toxicity (Fig. S4A). Translation inhibition of > 50% was only observed at 500 nM concentrations of gapmers 1, 2, 5 and 6 (Fig. S4B), which is at least 10-fold higher than the effective inhibition range for gapmers 1, 2 and 5. Overall, these results suggest that gapmers targeting the s2m element can inhibit viral replication in the model replicon system in a dosedependent, sequence-specific manner without causing significant cell toxicity.

### SARS-CoV-2 infection is inhibited by LNA gapmers targeting s2m

A high content screening (HCS) assay was developed to measure the effects of LNA gapmers on infection in Vero E6 cells infected with SARS-CoV-2. Figure 6A shows a graphical representation of the HCS assay workflow and an example representative microscopy image, with cells stained for N protein (488 nm signal). We tested gapmers 1-6, “all” and “scr” at 0.25, 0.5 and 1 μM concentrations, a no-gapmer control, and 10 μM remdesivir treated cells as a positive control. Our results indicate that gapmers 2 and 5 inhibit infection in a dose-dependent manner, reducing virus replication (measured through N protein expression) to 10.4 and 6.9%, respectively, of the non-treated control at 1 μM concentration, with cell viability at 72% and 83%, respectively. Furthermore, the inhibition levels are comparable to the 10 μM remdesivir control in this assay (Fig. 6B). Gapmer 1, 3, and 6 have a less profound effect (27%, 31%, and 17% of non-treated control levels at 1 μM), whereas gapmer 4 shows the highest toxicity levels besides inhibition to 13% of the non-treated control. In strong agreement with the astrovirus replicon-based results (Fig. 5D), gapmers 2 and 5 demonstrate the most promising results in SARS-CoV-2 inhibition assays (Fig. 6B), suggesting the suitability of these two gapmers for therapeutic development. Consistent with our previous results on other cell lines (Fig. S4A), the gapmers show no cytotoxic effect on Vero E6 cells in the absence of transfection reagent (<5%, Fig. 6C), providing further confidence for potential therapeutic gymnotic delivery (Fazil et al., 2016). These results are consistent with the astrovirus replicon-based approach and indicate that gapmers may have sufficient access to their RNA target in infected cells and that gapmers against the conserved s2m element may act as a viable anti-viral agent, warranting their further exploration as therapeutics.

**Figure 6.**
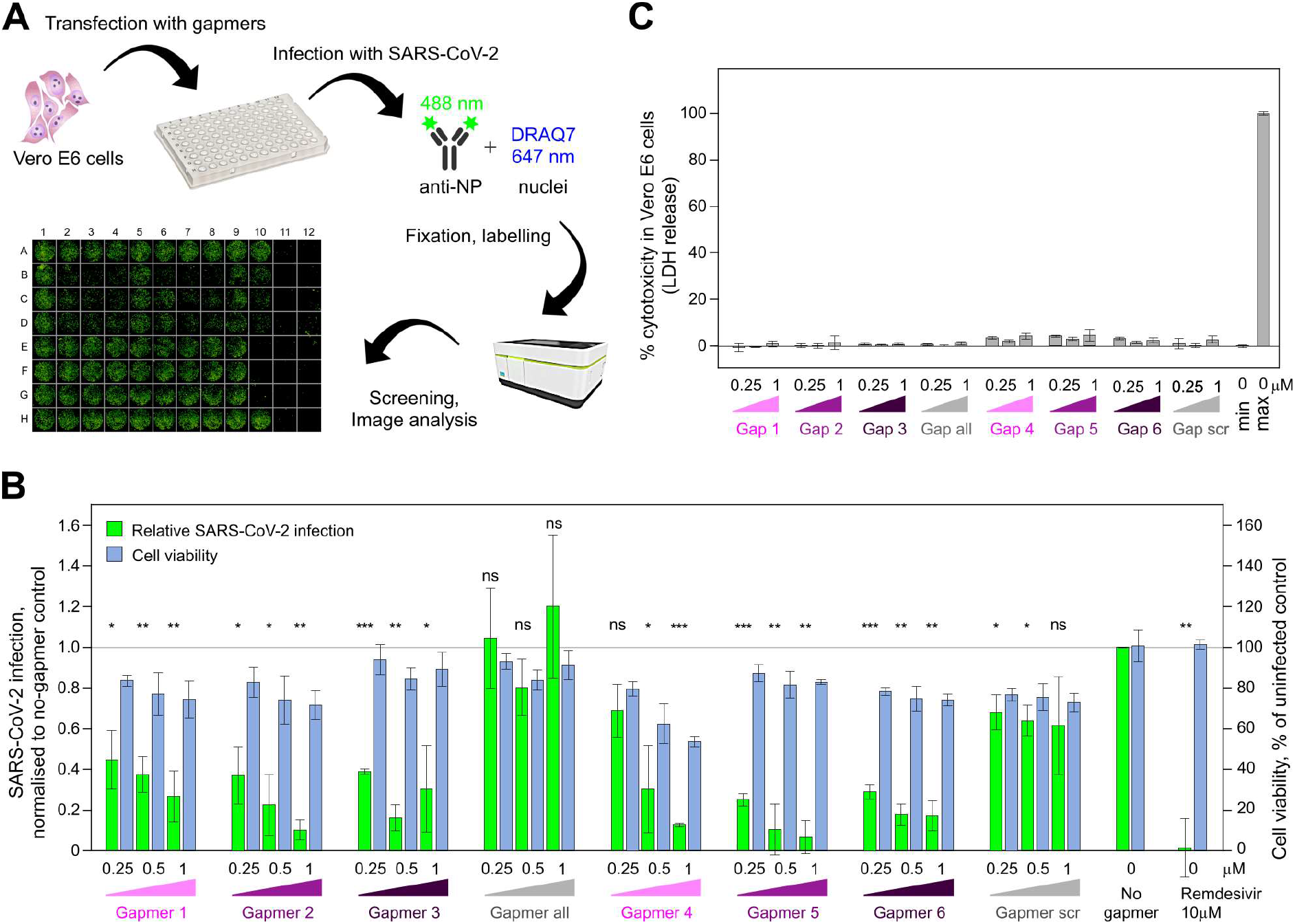
Inhibition of SARS-CoV-2 infection by gapmers targeting the s2m RNA element. (**A**) Graphic representation for the high content screening assay experiment workflow: transfection of Vero E6 cells with gapmers followed by infection with SARS-CoV-2, fixation of the plate, labelling and screening. A representative image from the immunofluorescence-based detection of SARS-CoV-2 infection of Vero E6 cells is shown below. (**B**) Vero E6 cells were transfected with gapmers 1-6 against the s2m element or control gapmers “all” and “scr” at 0.25, 0.5, and 1 μM final concentration and infected with SARS-CoV-2, fixed, labelled and analysed. Results are presented as mean ± s.d.; *n* = 3 biological replicates; signal normalised to a no-gapmer control. Cell viability was evaluated using the DRAQ7 signal normalised to mock treated wells. P-values are from two-tailed t-tests with separate variances (ns, *p* > 0.05; *, *p* ≤ 0.05; *p* ≤ 0.01; ***, *p* ≤ 0.001). (**C**) Toxicity assay for gapmer-treated Vero E6 cells in the absence of transfection reagent. Cells were treated with 0.25, 0.5, and 1 μM gapmers for 24 h. Supernatant was used to measure cell viability, calculated as the ratio of released to total lactate dehydrogenase (LDH) activity; “max” = maximum LDH measured for fully lysed cells. Full data and statistical analyses for B-C are provided in Supplementary Tables S3 and S4.

## DISCUSSION

We have confirmed that the highly conserved structural element s2m in the 3’ UTR of SARS-CoV-2 RNA is a site of vulnerability and a feasible candidate for ASO-based targeting by designing LNA-based gapmer ASOs effective against the s2m element *in vitro* and *in vivo.* We demonstrated physical gapmer-induced disruption of the RNA structure, consistent with successful gapmer-target base-pairing interactions, and RNase H-induced target degradation in *vitro*. Furthermore, we have demonstrated that the s2m element can direct gapmer-induced silencing of gene expression in GFP reporter assays in human cells, likely relying on the demonstrated enzymatic activity of endogenous RNase H in the nucleus and the cytoplasm (Liang et al., 2017).

We extended the observation by testing the ability of s2m-targeting gapmers to inhibit viral replication in an astrovirus-based replicon model system, a rapid functional assay that does not require access to higher containment level facilities. Viral replicon assays showed that gapmers against s2m inhibit viral replication in a sequence-specific, dose-dependent manner, down to sub-nanomolar range. These observations confirm that gapmers have significant potential as RNA replication inhibitors. Results from SARS-CoV-2 infection assays in cell culture with LNA gapmers targeting s2m are highly encouraging, with inhibition observed at the 0.25-1 μM gapmer concentration range. Two of the tested gapmers, 2 and 5, provided consistent inhibition results in both systems without increased cellular toxicity.

It is worth noting that coronaviruses produce double membrane vesicles inside infected host cells, thought to conceal the viral double-stranded RNA replicative intermediate from cellular defences (Hagemeijer et al., 2012; Knoops et al., 2008). These vesicles have been proposed to be formed through virus-induced manipulations of the membrane of endoplasmic reticulum (Blanchard and Roingeard, 2015). This compartmentalization of the SARS-CoV-2 genome in membranous bodies within the host cell’s cytoplasm (Klein et al., 2020) may reduce access of the ASOs to the s2m element or other viral genomic targets (although viral mRNAs, which all contain s2m, should still be accessible). In this regard, it is useful to consider that LNA ASOs have been conjugated with tocopherol and cholesterol for membrane association in the past (Benizri et al., 2019; Nishina et al., 2015), which could increase ASO ability to find its target. Improved membrane association may also boost gapmer entry into target cells. The primary targets for the SARS-CoV-2 infection are the ACE2 receptor expressing airway cells, with the virus infection gradually decreasing from the proximal to distal respiratory tract (Hou et al., 2020). These cells might be amenable to aerosol delivery (Drevinek et al., 2020), enabling highly targeted therapeutic administration.

When considering s2m as a target for virus inhibition, it is also worth noting that while it is a remarkably stable genomic element, conserved across multiple groups of single-stranded (+) sense RNA viruses, some mutations can arise over time. In the SARS-CoV-2 sequences from Covid-19 patient samples, some s2m polymorphisms have been detected at positions 15 and 31, predicted to destabilise the stem loop structure (Vahed et al, 2020; Yeh and Contreras, 2020). Using a combination of several gapmers together offers a potential strategy to guard against emerging resistance.

The results described here represent a promising start for further research into targeting conserved elements in single-stranded (+) sense RNA viruses, and support development of gapmers and related ASOs against the s2m element in particular. In the case of the SARS-CoV-2, our gapmer designs offer a strong starting point for further therapeutic development, which may include large scale optimisation and screening to maximise efficacy in cell culture and animal models, as well as chemical modifications for optimal delivery to target cells.

## ACKNOWLEDGEMENTS

AEF and VL are supported by Wellcome Trust (106207) and European Research Council (646891) grants. KJB, TD, KB and BFL are supported by a Wellcome Trust Investigator Award (200873/Z/16/Z) and TD by an Astra-Zeneca Studentship. X.Y and Y.D. are supported by a European Commission Horizon 2020 European Research Council (ERC) Starting Grant (680324). HJM and ND are supported by BBSRC (BBS/E/I/00007031 and BBS/E/I/00007037) grants. All cryoEM grids were prepared and cryoEM data collected at the BIOCEM facility, Department of Biochemistry, University of Cambridge. We thank Dimitri Y. Chirgadze, Steve Hardwick and Lee Cooper for assistance with data collection at the CryoEM Facility. We thank David LV Bauer for critical help with facilitating gapmer tests in virus infection assays. We thank Laura McCoy for the CR3009 antibody expression plasmids and Svend Kjaer at the Structural Biology Service Technology Platform at the Francis Crick Institute for preparation of the antibody. We thank Henrik Oerum, Alex Borodavka, Chris Oubridge and Ulrich Desselberger for invaluable advice and helpful discussions. We thank Dingquan Yu and Zhichao Miao for help with sequence analysis. We thank the support staff in our institutions for their invaluable help throughout the pandemic lockdown period. We dedicate this manuscript to the memory of our colleague Chris Oubridge.

## MATERIALS AND METHODS

### RNA preparation

3’ UTRs and the s2m 47-mer were prepared by *in vitro* transcription (IVT). Templates for IVT were generated either by PCR, using Phire Hotstart II polymerase (Thermo Fisher) according to manufacturer’s instructions, or by hybridising complementary DNA oligonucleotides (Sigma). Sequences for the PCR primers and the DNA oligonucleotides are given in Table 2. RNA generated by IVT was purified: IVT products were separated on polyacrylamide denaturing gel (National Diagnostics), relevant bands were excised using UV shadowing and electroeluted in TBE (Whatman Elutrap), then cleaned up using PureLinkTM RNA Microscale Kit (Invitrogen). RNA concentrations were estimated using UV absorbance (A260nm) and a calculated extinction coefficient.

### Gel-based RNA cleavage assay

Each gapmer was pre-incubated with target RNA in 1× RNase H buffer (Thermo Scientific) for 10 min at 37 °C. 2.5 U of RNase H (Thermo Scientific) was then added and the reaction incubated for 20 min at 37 °C. Reactions were quenched by adding an equal volume of proteinase K mix (0.5 mg/mL enzyme, 100 mM Tris-HCl pH 7.5, 150 mM NaCl, 12.5 mM EDTA, 1% w/v SDS) and incubating at 50 °C for 20 min. RNA was visualised on polyacrylamide 7.5 M urea gel (National Diagnostics) in TBE using SybrGold (Invitrogen).

### Chemical probing of the 3’ UTR

To probe RNA structure without gapmers, 5 pmol RNA (~350 ng) was dissolved in 9 μl nuclease-free water, and denatured at 95 °C for 90 s, then cooled on ice for 2 min. RNA was refolded by adding 10 μl of 2× SHAPE probing buffer (80 mM HEPES, pH 7.5, 200 mM KCl, 1 mM MgCl_2_) and incubating at 37 °C for 15 min. 2-methylnicotinic acid imidazolide (NAI) was added to a final concentration of 100 mM in 20 μl reaction; in control reactions, same volume of DMSO was added instead. Reactions were allowed to proceed at 37 °C for 5 min and then quenched by adding 10 μl of 2 M DTT. RNA was purified by loading quenched samples onto Micro Bio-spin Columns with Bio-Gel P-6 (BioRad), followed by ethanol precipitation. Purified RNA was then dissolved in 6 μl water, reverse-transcribed into cDNA and analysed by PAGE as described below.

To probe RNA structure with gapmers, different amounts of gapmers were added to refolded RNA, at molar ratios of 0.5×, 1× or 2×, and incubated at 37 °C for 10 min. The same volume of water was added for the no-gapmer control. After co-incubation, RNA was probed using NAI as described above. In addition, un-probed input RNA was also subjected to Sanger sequencing: RNA was dissolved in 5 μl water and supplemented with 1 μl of 10 mM corresponding ddNTP (Roche), then reverse-transcribed into cDNA and analysed by PAGE as described below.

### Reverse transcription and PAGE analysis of cDNA

6 μl of each RNA-containing sample was mixed with 1 μl of 5 μM Cy5-modified RT primer (sequence given in Table 2) and 0.5 μl of 10 mM dNTPs. The mixture was incubated at 95 °C for 3 min to denature the RNA and then cooled to 50 °C. RT reaction was started by adding 2 μl of 5× RT buffer (100 mM Tris pH 8.3, 500 mM LiCl, 15 mM MgCl_2_, 5 mM DTT) and 0.5 μl Superscript III enzyme (Invitrogen), mixing quickly with a pipette tip. Reaction was allowed to proceed at 50 °C for 20 min, then quenched by incubation at 85 °C for 10 min, which inactivates the enzyme. In order to degrade RNA and liberate the complementary cDNA, the reaction mix was then supplemented with 0.5 μl of 2 M NaOH and incubated at 95 °C for 10 min. The reaction was stopped by adding an equal volume of 2× stopping buffer (95% formaldehyde, 20 mM EDTA pH 8.0, 20 mM Tris-HCl pH 7.5, orange G dye) and incubating at 95 °C for 5 min. The resulting cDNA sample was cooled down to 65 °C and analysed on an 8% Acrylamide:Bis-Acrylamide-Urea gel by electrophoresis. The gel was imaged using Typhoon FLA 9000 Gel Imager (GE healthcare).

### Quantitative Gel Analysis and SHAPE Reactivity Calculation

Signal intensity of each band on the PAGE gel was detected using ImageQuant TL software and normalized to the total signal of the whole lane. Raw reactivity was generated by subtracting the signal of NAI channel from that of the DMSO control channel; reactivities with negative values were corrected to 0. SHAPE reactivity was generated following the 2/8 % normalization method (Low and Weeks, 2010). To calculate the differences in SHAPE signal in the presence and absence of gapmers, the lanes of NAI channels with 2× gapmer (+gapmer) or without gapmer (-gapmer) were used. The signal intensity of each band was normalized to the total signal of the whole lane. Differences were calculated by subtracting the signal of the +gapmer channel from that of the -gapmer channel and then rescaled to the signal of the - gapmer channel.

### Cryo electron microscopy

5 μM purified 3’ UTR in 10 mM Tris pH 7.6, 10 mM KCl, 10 mM NaCl was annealed by sequential incubation at 95 °C for 2 min, at 50 °C for 2 min, at 37 °C for 5 min and then at room temperature. The sample was supplemented with 1 mM MgCl_2_. 3 μl of the resultant mixture was applied to glow-discharged (EasiGlow Pelco) R2/2 Quantifoil grids (Quantifoil). Excess sample was blotted away with a FEI Vitrobot (IV) (100% humidity, 4 °C, blotting force 0, 3 s blot time) and the grids were vitrified in liquid ethane. The grids were screened with a 200 kV FEI Talos Arctica microscope with a Falcon III camera and a data set was collected on a 300 kV FEI Titan Krios microscope equipped with a Gatan K3 camera. Motion correction, ctf estimation and particle-picking were performed in Warp (Tegunov and Cramer, 2019) and 2D/3D alignments and averaging were carried out with cryoSPARC 2.15 (Punjani et al., 2017).

### Gapmer reporter assays

The s2m sequence or control scrambled sequence of s2m (s2m_scr) was inserted into the 3’ UTR of GFP in H6P plasmid using In-Fusion Cloning kit (TaKaRa) and verified by sequencing. HEK293ET, HeLa and A459 cells, as well as all stable cell lines, were grown in IMDM medium supplemented with 10% FCS at 37 °C in 5% CO_2_. All cell lines tested negative for mycoplasma. Constructs in H6P plasmids were used to produce recombinant lentivirus in HEK293ET cells for the stable expression of GFP reporter in mammalian cells from a constitutive SFFV promoter. HeLa and A549 stable cell lines were generated by lentiviral transduction with low MOI (>0.3) to ensure single genomic integrations and were selected for by drug resistance.

To examine the effect of gapmers on GFP expression, HeLa or A549 cells with a stably integrated GFP-s2m/s2m_scr reporter were seeded at 5 × 10^4^ cells per well in 24-well plates. The following day, cells were transfected with gapmer to achieve 20 nM final concentration using Lipofectamine RNAiMAX (Thermo Fisher Scientific). Flow cytometry analysis was performed 72 h after transfection. Cells were washed twice with warm PBS, detached with trypsin and resuspended in IMDM. All samples were analysed on a BD LSR ii flow cytometer, using the high throughput system (HTS), and at least 20,000 events were acquired for each sample. Data were analysed in FlowJo (v10.7.1). Main cell population was identified and gated on Forward and Side Scatter using the Auto Gate tool and plotted as a histogram to visualise GFP intensity of cells (measured using B-525 detector).

### Virus replicon assays

HEK293T cells (ATCC) were maintained at 37 °C in DMEM supplemented with 10% fetal bovine serum (FBS), 1 mM L-glutamine, and antibiotics. Huh7.5.1 cells (obtained from Apath, Brooklyn, NY; Zhong et al., 2005) were maintained in the same media supplemented with nonessential amino acids. All cells were mycoplasma tested (MycoAlert™ PLUS Assay, Lonza); Huh7.5.1 cells were also tested by deep sequencing.

The HAstV1 replicon system is based on the HAstV1 infectious clone, where the virus genome is left intact up to the end of ORFX, then followed by a foot-and-mouth disease virus 2A sequence and a *Renilla* luciferase (RLuc) sequence fused in the ORF2 reading frame, followed by the last 624 nt of the virus genome and a poly-A tail (Fig. 4A, GenBank accession number MN030571; Lulla and Firth, 2020). The s2m mutations were introduced using site-directed mutagenesis and all constructs were confirmed by sequencing. The resulting plasmids were linearized with *Xho*I prior to T7 RNA transcription.

Huh7.5.1 and HEK293T cells were transfected in triplicate with Lipofectamine 2000 reagent (Invitrogen), using the protocol in which suspended cells are added directly to the RNA complexes in 96-well plates. For each transfection, 100 ng replicon, 10 ng firefly luciferase-encoding purified T7 RNA (RNA Clean and Concentrator, Zymo research), the indicated amount of gapmers, and 0.3 μl Lipofectamine 2000 in 20 μl Opti-Mem (Gibco) supplemented with RNaseOUT (Invitrogen; diluted 1:1,000 in Opti-Mem) were added to each well containing 5 × 10^4^ cells in 100 μl DMEM supplemented with 5% FBS, and incubated at 37 °C for 12 h (Huh7.5.1) or 18 h (HEK293T). Firefly and Renilla luciferase activities were determined using the Dual Luciferase Stop & Glo Reporter Assay System (Promega). Replicon activity was calculated as the ratio of Renilla (subgenomic reporter) to Firefly (cap-dependent translation, loading control), normalized by the same ratio for the control replicon, as indicated for each experiment. Three independent experiments, each in triplicate, were performed to confirm reproducibility of results.

### Cytotoxicity assays

The analysis of gapmer cellular cytotoxicity in HEK293T, Huh7.5.1, and Vero E6 cells was performed using the CyQUANT LDH cytotoxicity assay (Thermo Scientific). Leaked cytoplasmic enzyme LDH in cell culture supernatants was quantified after enzymatic conversion, and absorbance was measured at 490 nm in a 96-well plate reader according to the manufacturer’s instructions.

### SARS-CoV-2production and infection

Vero E6 cells obtained from Oliver Schwarz (Institute Pasteur, Paris) were maintained in DMEM modified with high glucose, L-glutamine, phenol red and sodium pyruvate (ThermoFisher, #41966-029) supplemented with 10% FCS at 5% CO_2_. SARS-CoV-2 strain BetaCoV/England/02/2020 (obtained from Public Health England) was propagated at 37°C on Vero E6 cells in DMEM supplemented with 10% FCS at 37 °C. The titer was determined by plaque assay as follows: confluent monolayers of VeroE6 cells where grown on 6-well plates and incubated with 200 μl of a 10-fold serial dilution of virus stock in DMEM supplemented with 10% FCS for 1 h at room temperature. These cells were then overlaid with 0.5× DMEM supplemented with 1% FCS and 1.2% Avicel (BMC Biopolymers, Belgium). After 4 days incubation at 37 °C, cells were fixed with 4% formaldehyde in PBS followed by staining with 0.1% toluidine blue (Sigma, #89640). The titer was calculated as plaque forming units (PFU) per ml.

### SARS-CoV-2 infection assay

Vero E6 cells (NIBC, UK) were grown in DMEM (containing 10 % FBS) at 37 °C and 5 % CO_2_ in 96 well imaging plates (Greiner 655090). Cells were transfected with individual gapmers at 0.25, 0.5 or 1 μM final concentration using Lipofectamine 2000 (Thermo Fisher), according to manufacturer’s instructions. At 6 h post transfection, the media was replaced and the cells were infected or mock infected with SARS-CoV-2 at a multiplicity of infection 0.5 PFU/cell. At 22 h post-infection, cells were fixed, permeabilised and stained for SARS-CoV-2 N protein using Alexa488-labelled-CR3009 antibody (van den Brink et al., 2005) and for nuclei (DRAQ7, 647 nm wavelength). The plate was imaged using the high content screening microscope Opera Phenix from Perkin Elmer with a 5× lens. The Opera Phenix-associated software Harmony was used to delineate the whole well area and to determine the total intensities of the Alexa488/N protein and DRAQ7/DNA signals per said whole well area during image acquisition. The background subtracted Alexa488/N intensities were normalised to vehicle treated samples. Toxicity was evaluated using the DRAQ7/DNA signal normalised to mock treated wells and LDH release-based cytotoxicity assay.

## SUPPLEMENTARY FIGURES

**Figure S1, related to Figure 1.**
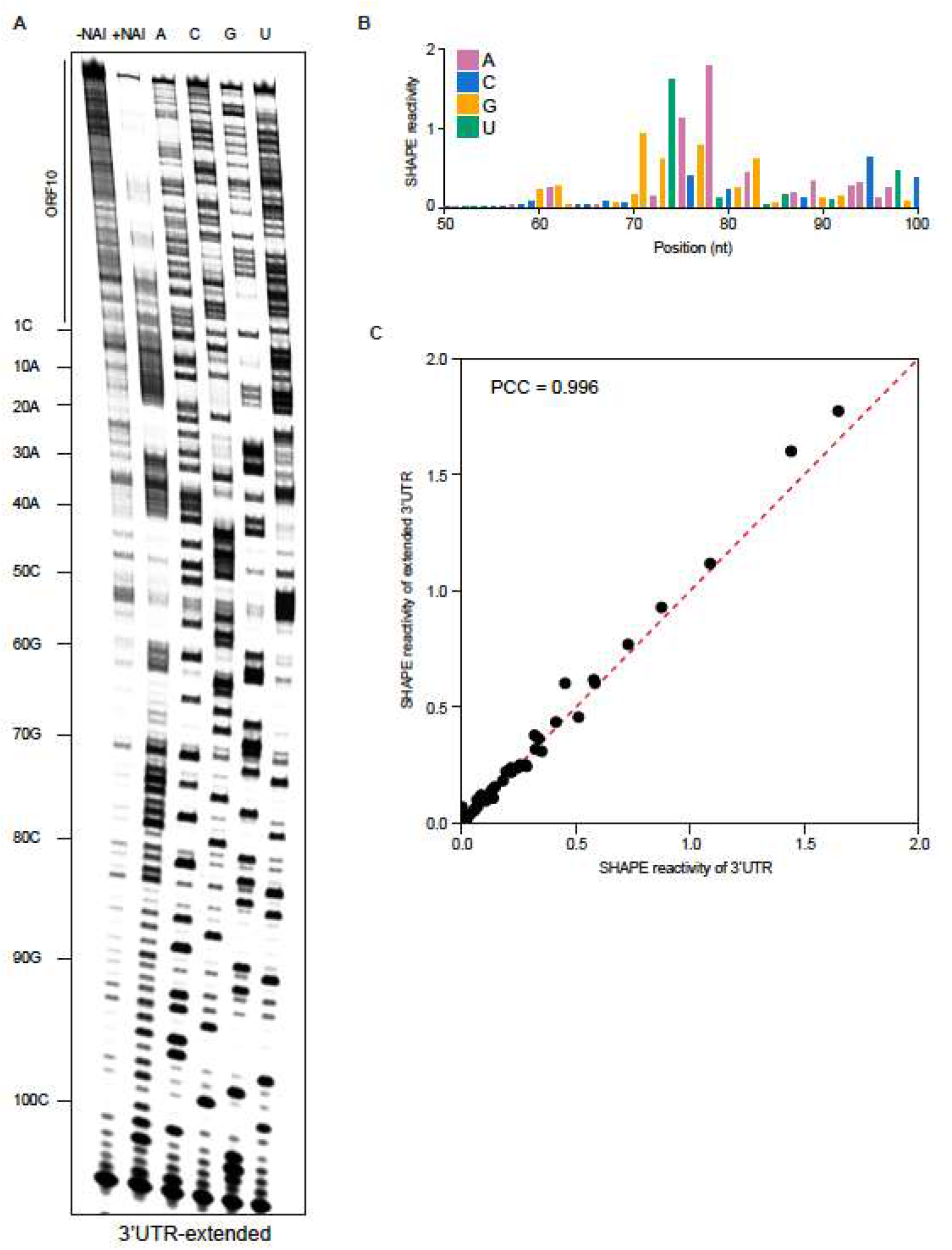
SHAPE analysis of the extended 3’ UTR of SARS-CoV-2. (**A**) Chemical probing of the extended 3’ UTR of SARS-CoV-2 (including ORF10 and the region immediately upstream of it). RNA was denatured and refolded in the presence of 100 mM K^+^ and 0.5 mM Mg^2+^, then incubated with NAI (+NAI channel) or DMSO control (- NAI channel). NAI modification was detected by reverse transcription stalling and gel-based analysis. Sequencing lanes were generated by adding ddT (for A), ddG (for C), ddC (for G) and ddA (for U) when performing reverse transcription. (**B**) Quantification of SHAPE signal on at the s2m element and flanking regions. Calculation was based on the gel in Fig. S1A, by subtracting the signal of the +NAI lane from that of the -NAI lane. (**C**) Agreement of SHAPE signal on s2m structure in the 3’ UTR and the extended 3’ UTR of SARS-CoV-2. Pearson correlation coefficient was calculated based on the SHAPE signals shown in Fig. 1C and Fig. S1B.

**Figure S2, related to Figure 4.**
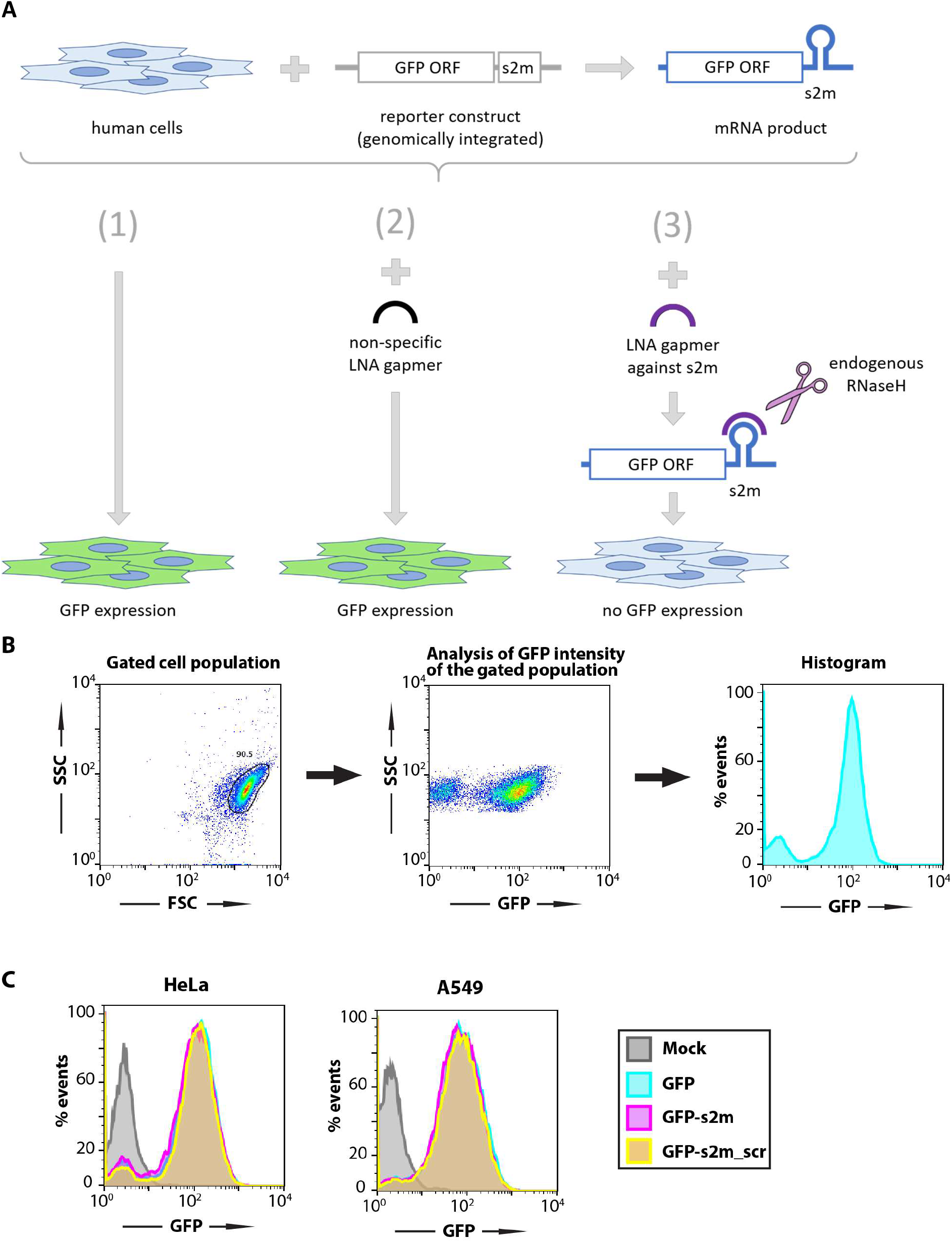
s2m element does not influence RNA translation in the reporter assay in human cells. (**A**) Schematic of the GFP reporter assay. (**B**) Gating and data visualisation strategy of the reporter assay data. Main cell population was identified and gated on Forward and Side Scatter using the Auto Gate tool and plotted as a histogram to visualise GFP intensity of cells. (**C**) HeLa and A549 cell lines containing a genomic insertion of a GFP reporter construct without additional insertion in its 3’ UTR (GFP), with the s2m sequence in its 3’ UTR (GFP-s2m) or with a scrambled sequence insertion in its 3’ UTR (GFP-s2m_scr) were analysed by flow cytometry. Data are representative of two independent experiments.

**Figure S3, related to Figure 4.**
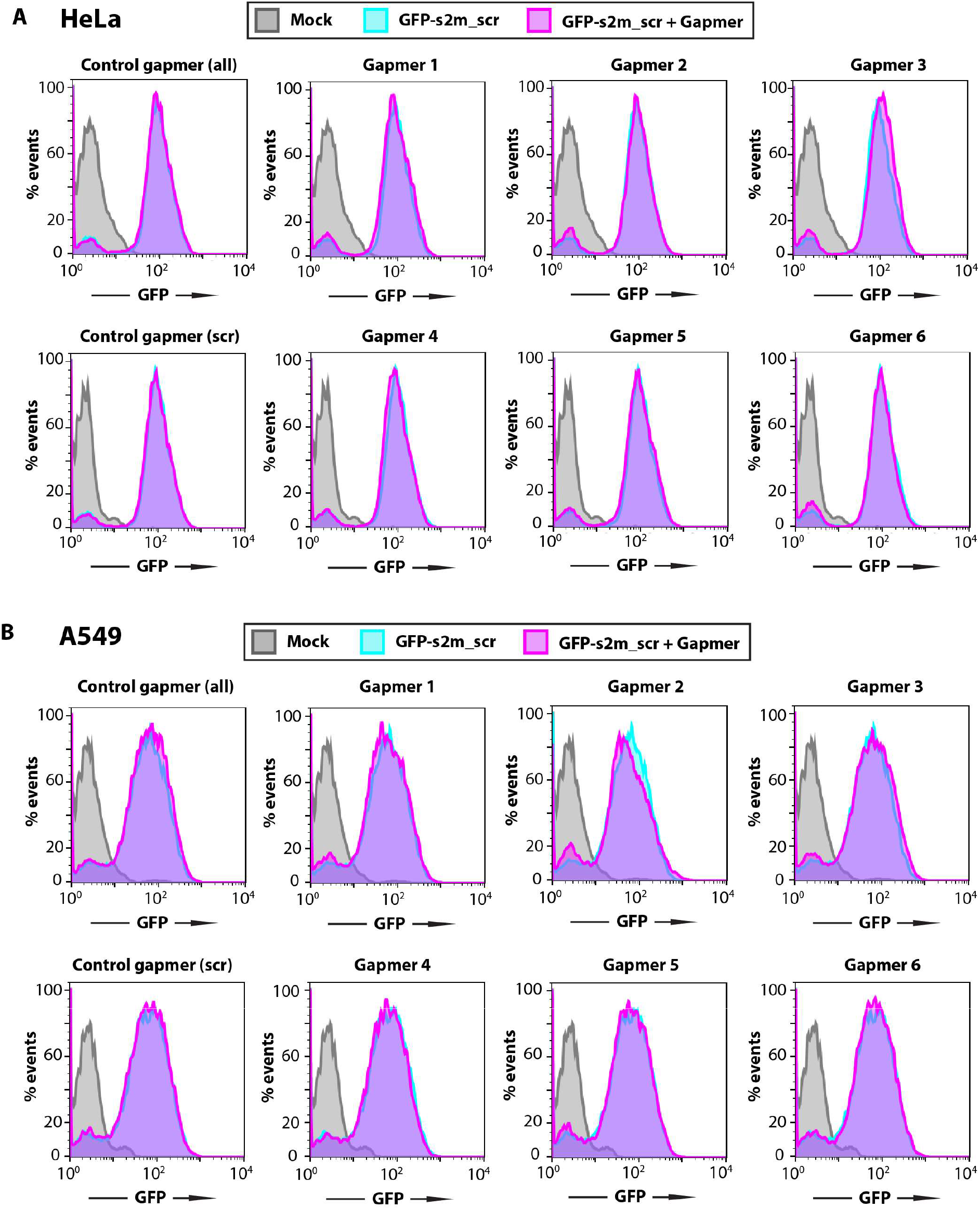
Gapmer silencing effect relies of sequence-specific gapmer-target interaction. HeLa (**A**) and A549 (**B**) cell lines containing a control genomic insertion of a GFP reporter construct with a scrambled sequence inserted in its 3’ UTR (GFP-s2m_scr) were transfected with 20 nM of the indicated gapmers and analysed 72 h post-transfection by flow cytometry. The control cell lines containing GFP reporter with a scrambled sequence insertion in the 3’ UTR show no appreciable change in fluorescence upon treatment with the gapmers targeted against the s2m element. Data are representative of three independent experiments.

**Figure S4, related to Figure 5.**
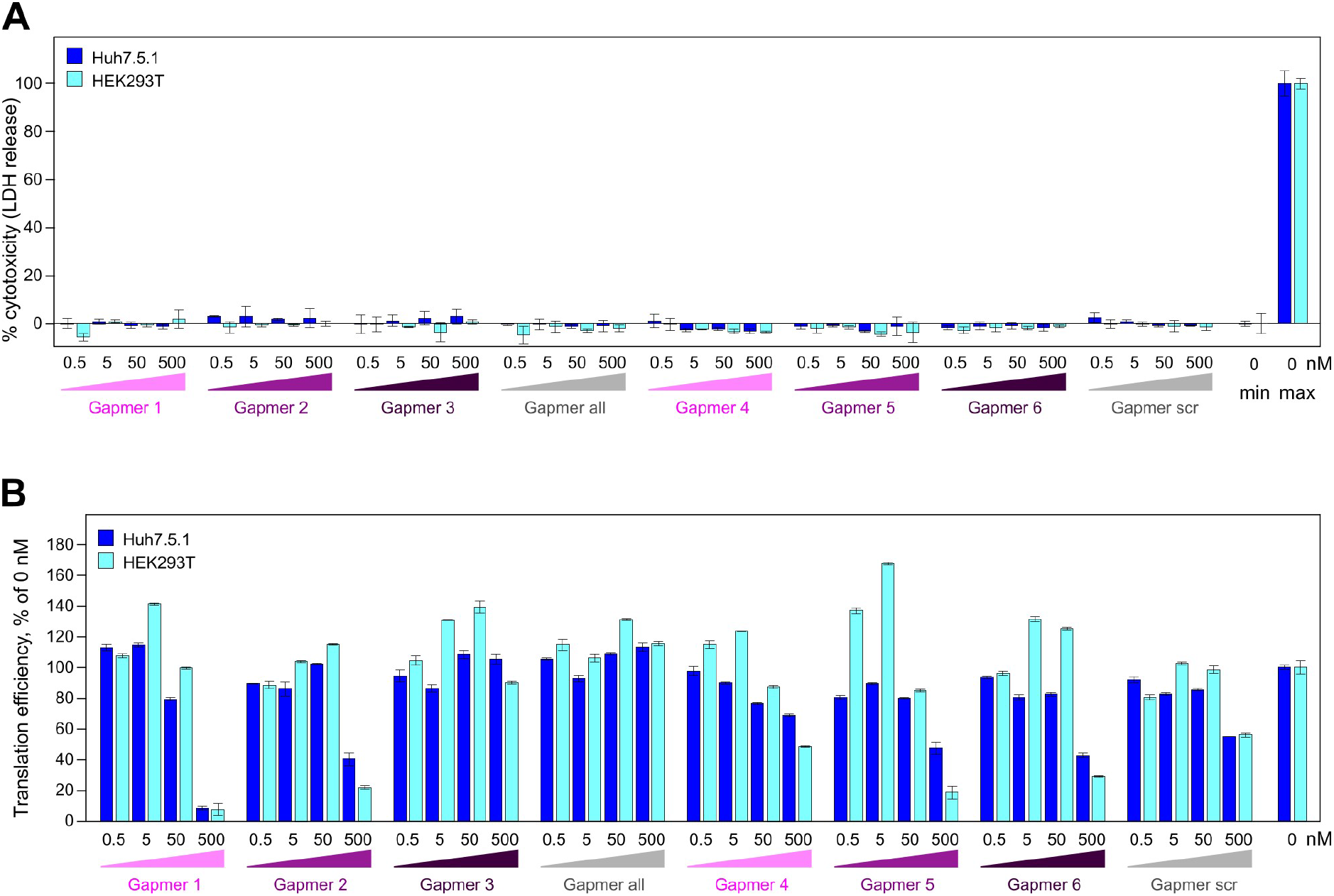
Testing the cytotoxic and off-target effects of gapmers. (**A**) Toxicity assay for gapmer-treated cells. Cells were treated with 0.5-500 nM gapmers for 24 h. Supernatant was used to measure cell viability, calculated as the ratio of released to total lactate dehydrogenase (LDH) activity; “max” = maximum LDH measured for fully lysed cells. (**B**) The effect on translation measured as a readout of capped T7 RNA encoding firefly luciferase at 0.5-500 nM gapmer concentration, normalized to untreated cells. All data are presented as mean ± s.d.; *n* = 3 biologically independent experiments.

